# The tubovesicular network in *Plasmodium vivax* liver-stage hypnozoites and schizonts associates with host aquaporin 3

**DOI:** 10.1101/2020.12.09.417949

**Authors:** Kayla Sylvester, Steven P. Maher, Dora Posfai, Michael K. Tran, McKenna C. Crawford, Amélie Vantaux, Benoît Witkowski, Dennis E. Kyle, Emily R. Derbyshire

## Abstract

The apicomplexan *Plasmodium* parasites replicate in the liver before causing malaria. *P. vivax* can also persist in the liver as dormant hypnozoites and cause relapses upon activation. The host water and solute channel aquaporin-3 (AQP3) has been shown to localize to the parasitophorous vacuole membrane (PVM) of *P. vivax* hypnozoites and liver schizonts, along with other *Plasmodium* species and stages. In this study, we use high-resolution microscopy to characterize temporal changes of the tubovesicular network (TVN), a PVM-derived network within the host cytosol, during *P. vivax* liver-stage infection. We demonstrate an unexpected role for the TVN in hypnozoites and reveal AQP3 associates with TVN-derived vesicles and extended membrane features. We further show AQP3 recruitment to *Toxoplasma gondii*. Our results highlight dynamic host-parasite interactions that occur in both dormant and replicating liver-stage *P. vivax* forms and implicate AQP3 function during this time. Together, these findings enhance our understanding of AQP3 in apicomplexan infection.

## Introduction

Apicomplexans constitute a large group of parasitic protozoans that cause diseases, including the five species of *Plasmodium* that cause human malaria. Human infection by *Plasmodium* begins when a female *Anopheles* mosquito deposits sporozoites into the bloodstream which then migrate to the liver. Once in a hepatocyte, sporozoites transform and rapidly replicate to yield tens of thousands of merozoites from a single schizont (Ejigiri and Sinnis, 2009; Frischknecht et al., 2004; Ploemen et al., 2009; Prudêncio et al., 2006). This stage is asymptomatic but a prerequisite to the infection of red blood cells (RBCs) that leads to disease. Many species of *Plasmodium*, including *P. falciparum* and *P. berghei*, follow this path, while others such as *P. vivax* can differentiate in hepatocytes to either schizonts or hypnozoites, a biologically quiescent form (Krotoski et al., 1982). Dormant hypnozoites cause recurrent blood-stage infections (relapses) upon activation, effectively sustaining repeated blood infection and furthering transmission (Krotoski et al., 1982; Mikolajczak et al., 2015). *P. vivax* hypnozoites are insensitive to most antimalarials except 8-aminoquinolines, which are contraindicated in populations with G6PD deficiency, highlighting the need for new anti-hypnozoite agents (Howes et al., 2012; Lu and Derbyshire, 2020). Unfortunately, efforts to identify compounds capable of inhibiting hypnozoites are hampered by our limited understanding of the molecular pathways that enable parasite survival and activation during this stage.

To successfully develop and replicate within their host, *Plasmodium* resides within a membrane-bound compartment formed during invasion termed the parasitophorous vacuolar membrane (PVM) (Meis et al., 1983). Derived from the host membrane itself, the PVM serves as the host-pathogen interface and dynamically changes to facilitate development by enabling nutrient acquisition and protecting the parasite from apoptosis (Kaushansky et al., 2013; Meireles et al., 2017; Sá E Cunha et al., 2017; Van De Sand et al., 2005; Van Dijk et al., 2005). For example, the PVM diameter can expand beyond that of the pre-invasion hepatocyte membrane diameter and can recruit both host and parasite proteins. The precise composition of this membrane remains unknown, but several studies have revealed it changes minutes to hours after liver invasion when parasite proteins, including up-regulated in infective sporozoites gene 4 (UIS4), are translocated to the PVM (Kaushansky and Kappe, 2015; Mueller et al., 2005; Prado et al., 2015; Sá E Cunha et al., 2017; Schnider et al., 2018). While fewer host proteins are known to associate with the PVM after invasion, examples of protein recruitment crucial for host defense and parasite development have been reported (Grützke et al., 2014; LaMonte et al., 2019; Niklaus et al., 2019; Posfai et al., 2020, 2018; Prado et al., 2015; Raphemot et al., 2019; Thieleke-Matos et al., 2016; Wacker et al., 2017). In particular, we have demonstrated that the host water and solute channel aquaporin 3 (AQP3) is upregulated after *P. berghei* infection of liver cells, is critical for parasite development, and associates with the PVM (Posfai et al., 2018). Importantly, we have established the recruitment of AQP3 in multiple *Plasmodium* species and stages including liver-stage *P. berghei*, blood-stage *P. falciparum*, blood-stage *P. vivax*, and liver-stage *P. vivax* (schizonts and hypnozoites) (Posfai et al., 2020, 2018). While the reduced intrahepatic *P. berghei* size observed with genetic or chemical inhibition of AQP3 supports an essential role in *Plasmodium* development, its molecular function during infection has yet to be resolved.

A key component of the host-parasite interactions in the *Plasmodium* liver stage is the physical alteration of the PVM. Several elegant studies have shown that the PVM can extend into the host hepatocyte with a highly dynamic membranous system, the tubovesicular network (TVN) (Agop-Nersesian et al., 2018, 2017; Grützke et al., 2014; Niklaus et al., 2019). In the *P. berghei* liver-stage, this system consists of elongated membrane clusters, tubules, and vesicles that move to and from the PVM (Grützke et al., 2014). The TVN expansion into the host cytosol is proposed to aid in nutrient acquisition and immune evasion, imparting key survival strategies to the parasite (Agop-Nersesian et al., 2018, 2017; Grützke et al., 2014; Niklaus et al., 2019). While UIS4 is an established parasite-derived *P. berghei* TVN marker, the only host proteins reported to associate with TVN features are LC3, LAMP1, p62 and CD63 (Agop-Nersesian et al., 2018, 2017; Grützke et al., 2014; Niklaus et al., 2019). There has been no evidence of a protein responsible for nutrient uptake to be associated with the TVN. In the blood-stage, the TVN is complemented by an unusual protein trafficking system that involves long-known parasite-derived membranous structures in the RBC cytosol, termed Maurer’s clefts in *P. falciparum* and Schüffner’s dots in *P. vivax* (Akinyi et al., 2012; Alkawa et al., 1975; Mundwiler-Pachlatko and Beck, 2013; Sakaguchi et al., 2016; Schüffner, 1899; Spycher et al., 2006; Tamez et al., 2008; Tokumasu et al., 2014; Wickert et al., 2004; Wickert and Krohne, 2007). Notably, these structures have not been found in infected hepatocytes. Additionally, the TVN has not yet been reported in liver-stage *P. vivax*, likely due to the technical difficulties associated with studying parasites during this stage.

Recent advances in *P. vivax* model systems that can support liver-stage schizont and hypnozoite development have been instrumental to setting the stage for anti-hypnozoite drug screens and aiding molecular studies to understand parasite biology (Antonova-Koch et al., 2018; Gural et al., 2018; Maher et al., 2020; March et al., 2013; Mikolajczak et al., 2015; Roth et al., 2018). Here, we analyzed *P. vivax*-infected primary human hepatocytes (PHH) to discover and characterize TVN features throughout infection (Roth et al., 2018). Interestingly, we observe that TVN features are more abundant in dormant hypnozoites than in replicating schizonts. We also found that AQP3 co-localizes with TVN vesicles and extended membrane clusters at certain time points in *P. vivax* hypnozoites as well as liver-stage *P. vivax* and *P. berghei* schizonts, implicating a role in solute transport or evasion of host defense mechanisms. Additionally, we observe AQP3 localization to the apicomplexan parasite *Toxoplasma gondii* and upregulation of *AQP3* expression after parasite infection of hepatocytes. Together, our work suggests a possibly conserved role for AQP3 in apicomplexans to aid in development through nutrient acquisition or protection from host defenses.

## Results

### *P. vivax* TVN characterization

The *P. berghei* liver-stage TVN is known to be important for nutrient acquisition and immune evasion, but this network has not yet been characterized in the *P. vivax* liver-stages. *P. berghei* requires approximately 2 days to mature from a sporozoite to a schizont in the liver (Figure 1—figure supplement 1), while *P. vivax* schizont maturation occurs over 8-10 days in vitro. *P. vivax* also has an alternative morphological and physiological route that arrests development to create a quiescent hypnozoite that can activate days or months after infection (Figure 1A). To examine the possibility of TVN formation in *P. vivax* schizonts or hypnozoites, we used high-resolution confocal microscopy to obtain images of *P. vivax*-infected PHH 2–10 days post-infection (dpi). Parasites were stained with a recombinant anti-*Plasmodium* UIS4 (rUIS4) antibody (Schafer et al., 2018) to visualize the PVM/TVN and DAPI to evaluate number of parasite nuclei. This analysis showed that TVN features were present throughout PHH infection with *P. vivax*, with a higher percentage of exoerythrocytic forms (EEFs) positive for TVN features when observed on days 5 and 6 post-infection (Figure 1B). While the schizont and hypnozoite populations are of similar sizes and are difficult to distinguish up to 5-6 dpi, our analysis suggested TVN features were more abundant in smaller sized parasites when observed at 8 dpi. To explore this observation, we performed two classifications. First, we measured the area of each EEF and divided them into one of two categories: not exhibiting TVN features or exhibiting features at 8 dpi (Figure 1C). This analysis revealed that the vast majority of *P. vivax* EEFs displaying TVN features were relatively small at 8 dpi (Figure 1D). Second, to definitively characterize EEF populations as hypnozoites and schizonts, we quantified the number of parasite nuclei and growth area of each EEF when imaged at high resolution (63x magnification). A uninucleated parasites indicates a non-replicating dormant hypnozoite, while a multinucleated parasite indicates a schizont or activated hypnozoite, as either would be actively replicating (Mikolajczak et al., 2015). We observed all multinucleated EEFs as having an area ≥75 cm^2^, thus we established a strict size cutoff for hypnozoites as having a growth area <60 µm^2^ (Figure 1—figure supplement 2A) and for schizonts as having a growth area above >200 µm^2^(Figure 1—figure supplement 2B). While few net 8 dpi EEFs were noted with a growth area between 60 µm^2^ and 200 µm^2^, these forms were excluded from classification of hypnozoite or schizont to ensure no ambiguity between the two forms. This more precise analysis of the small parasite population revealed the vast majority of the EEFs displaying TVN features were classified as hypnozoites. Using these metrics, we found that 32% of hypnozoites had one or more TVN features at 8 dpi (Figure 1E), while no schizonts displayed features among the 529 EEFs analyzed on this dpi.

**Figure 1.**
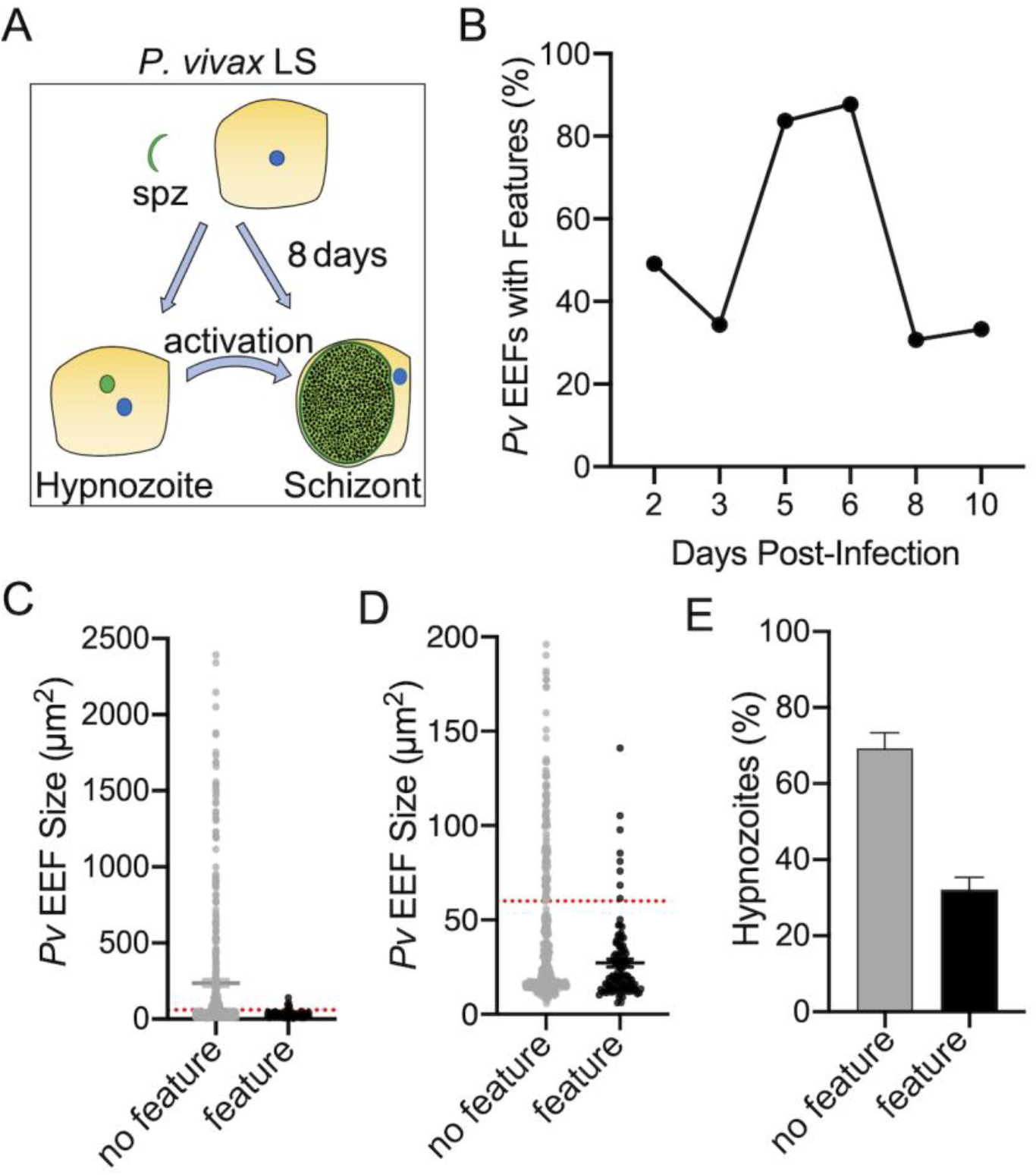
*P. vivax*-infected PHH contain TVN features. (A) Schematic representing *P. vivax* LS infection. (B) Percentage of *P. vivax* EEFs that contain features (black) throughout infection of PHH (2–10 dpi). (C) Relative size of *P. vivax* EEFs containing no TVN features (grey) or TVN feature/s (black). (D) Relative size of *P. vivax* EEFs (from C), displaying smaller scale, containing no TVN features (grey) or TVN feature/s (black). Red dotted line indicates cut off for hypnozoites (60 µm^2^). (E) Percentage of *P. vivax* hypnozoites exhibiting no TVN features (grey bar) or TVN feature/s (black bar). (C-E) Assessed at 8 dpi (3 independent wells assessed with >200 EEFs, n=2).

Through our analysis, we observed that most *P. vivax* liver-stage EEFs displaying a TVN had multiple features (2–3). To further examine these features and the relationship with *P. vivax* hypnozoites and schizonts, we classified *P. vivax* TVN features as extended membrane clusters (mem. cluster), tubules, and vesicles (Figure 2A), similar to a previous liver-stage TVN *P. berghei* study (Grützke et al., 2014). This analysis revealed that TVN-derived vesicles in the host cytosol were most prominent in cells containing hypnozoites at 8 dpi, present within 30% of the population (Figure 2B). Extended membrane clusters and tubules were present in ∼15% of the hypnozoite population. Throughout infection, the *P. vivax* TVN includes multiple features, where the majority of EEFs displaying extended membrane clusters or tubules also displayed TVN-derived vesicles (Figure 2—figure supplement 1A). These TVN-derived vesicles are sometimes seen close to the EEF, but also can be found at distal parts of the cell.

**Figure 2.**
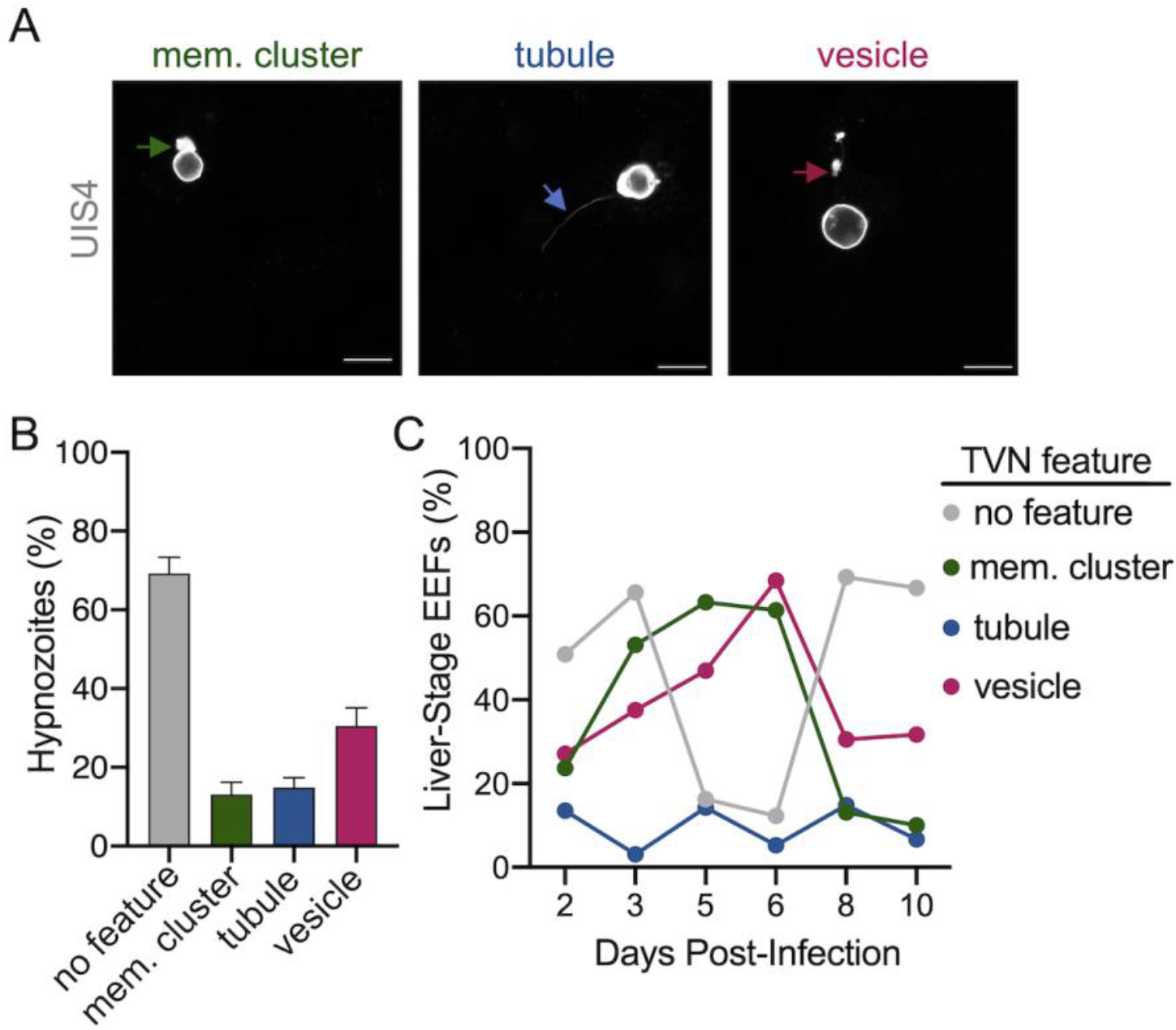
Characterization o*f P. vivax* TVN throughout infection. (A) Examples of TVN features including extended membrane cluster (mem. cluster, green arrow), tubule (blue arrow) and vesicles (magenta arrow) in *P. vivax*-infected PHH at 8 dpi. Cells stained with anti-*Pv*UIS4 (grey). Scale bar is 10 µm. (B) Characterization of *P. vivax* hypnozoites at 8 dpi containing no feature or features. Percentage of each characteristic including extended membrane cluster (mem. cluster, green bar), tubule (blue bar), and vesicle (magenta bar) is indicated. Grey bar indicates percentage of hypnozoites with no features (3 independent wells assessed with >200 EEFs, n=2). (C) Percentage of total *P. vivax* EEFs with indicated TVN characteristic days 2–10 post-infection of PHH (n>30 EEFs each day).

We hypothesized that TVN features may vary depending on the parasite’s developmental stage. Therefore, we took a longitudinal approach to our analysis and evaluated TVN features every day of a 10-day infection of PHH with *P. vivax*. Since hypnozoites and schizonts cannot be reliably distinguished by size before 5-6 dpi, EEFs were not separated into hypnozoites or schizonts. We discovered variation in the prevalence of different TVN features throughout infection (Figure 2C). In *P. berghei*, TVN features are more prevalent at early times of hepatocyte infection, appearing within 30 min and being most abundant between 6–24 hours post-infection (hpi) (Grützke et al., 2014). Here, we observe that early *P. vivax* EEF stages (2–3 dpi), display few TVN features, but a marked increase in extended membrane clusters and TVN-derived vesicles occurs from 2 to 6 dpi. Based on this analysis, 3–5 and 6–8 dpi are particularly dynamic times where the greatest changes in TVN abundance and composition were observed. For example, a remarkable decrease in extended membrane clusters is observed between 6 to 8 dpi, such that less are present at 8 dpi than at the early time points (i.e. 2 dpi). While a decrease in TVN-derived vesicles was also observed between 6 to 8 dpi, the prevalence of this feature within cells remained high compared to that of extended membrane clusters. TVN-derived vesicles were maintained in >20% of EEFs throughout the course of infection. Relatively less change in tubule abundance was observed throughout intrahepatic *P. vivax* development. These tubules vary in length and, interestingly, we have observed some instances of tubule association with the host nuclei (Figure 2—figure supplement 1B). On days 8–10 post-infection, when schizonts can be distinguished from hypnozoites, we evaluated TVN features as a function of size. Though we did not observe TVN features in schizonts at 8 dpi, we observed a TVN positive population of schizonts at days 9 and 10 (Figure 2—figure supplement 2A). Hypnozoites also displayed TVN features at 9 and 10 dpi (Figure 2—figure supplement 2B).

### AQP3 is associated with the *Plasmodium* TVN

The precise compositions of the *Plasmodium* liver-stage PVM and TVN remain to be resolved. We recently established that host AQP3 is recruited to the PVM of *P. berghei* (Posfai et al., 2018) and *P. vivax* EEFs, including both developing schizonts and dormant hypnozoites (Posfai et al., 2020, 2018). To better understand AQP3 recruitment to these different forms, we visualized *P. berghei*-infected HuH7 cells and *P. vivax*-infected PHH after staining with DAPI, anti-rUIS4 and anti-human AQP3 antibodies (Posfai et al., 2020, 2018). We observed that AQP3 co-localizes to some, but not all TVN features in *P. berghei* and *P. vivax* EEFs at 2 and 8 dpi, respectively (Figure 3A and Figure 3—figure supplement 1). To further investigate AQP3 recruitment to the TVN, we analyzed *P. vivax* EEFs days 2–10 post-infection (Figure 3B). At early time points (2–3 dpi), AQP3 is not observed in every *P. vivax* EEF (Posfai et al., 2020); however, it was highly recruited to extended membrane clusters and vesicles when present. For example, when an EEF exhibited both AQP3 and extended membrane TVN features at 2 dpi, they were observed together in every EEF examined in our analysis. Likewise, AQP3 was highly correlated with TVN-derived vesicles at 3 dpi. Notably, we observed no AQP3 recruitment to TVN features in EEFs that lacked protein recruitment to the PVM. Further, we found no instance of AQP3 co-localizing with *P. vivax* TVN tubules at any day post infection despite looking in multiple planes (z-stacks).

**Figure 3.**
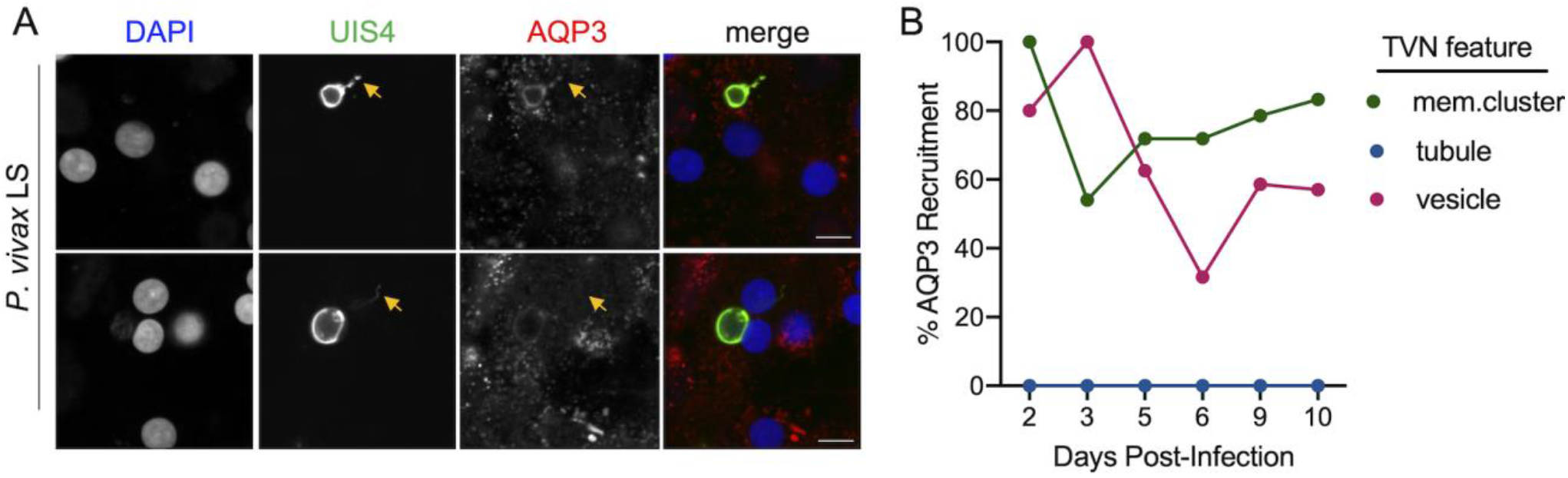
Localization and recruitment of AQP3 to TVN features during *P. vivax* infection. (A) Representative confocal images of *P. vivax* EEFs exhibiting TVN features, stained with DAPI (blue), anti-*Pv*UIS4 (green), and anti-*Hs*AQP3 (red) at 8 dpi. *P. vivax*-infected PHH show localization (top) and no localization (bottom) of AQP3 to specific TVN features (yellow arrows). Scale bar is 10 µm. (B) Relative percentage of TVN features in *P. vivax*-infected PHH exhibiting AQP3 recruitment (2_–_10 dpi, n>30 EEFs per dpi).

### AQP3 is recruited during *T. gondii* infection

We sought to more broadly examine the role of AQP3 in apicomplexan parasites. In particular, *T. gondii*, the causative agent of toxoplasmosis, is considered a model apicomplexan organism due to its ability to infect almost any multinucleate cell with a high infection rate (Black and Boothroyd, 2000). This intracellular parasite also resides in a parasitophorous vacuole which creates a barrier between the host cytoplasm and the growing parasite. We evaluated the recruitment and gene expression of AQP3 after *T. gondii* Pru A7 infection of various host cells. In *T. gondii*-infected HuH7 cells, we observed recruitment of AQP3 to parasites at both 24 hpi and 48 hpi (Figure 4A). At 24 hpi, AQP3 is observed in ∼10% of infected cells and is associated with the parasite plasma membrane (PPM). By 48 hpi, AQP3 recruitment is observed in >50% of infected HuH7 cells and it is most commonly observed surrounding tachyzoite rosettes. At this time AQP3 is also observed at the residual body of some but not all rosettes (Figure 4—figure supplement 1). In addition to *T. gondii* Pru A7-infected HuH7 cells, AQP3 recruitment is observed in *T. gondii* Pru A7-infected Vero cells at 24 and 48 hpi (Figure 4—figure supplement 2A) and in *T. gondii* RH-infected HuH7 cells at 48 hpi (Figure 4—figure supplement 2B). *T. gondii* Pru A7 is a type II strain while *T. gondii* RH is type I. These results demonstrate AQP3 recruitment to various *T. gondii* strains and in different mammalian cell lines.

**Figure 4.**
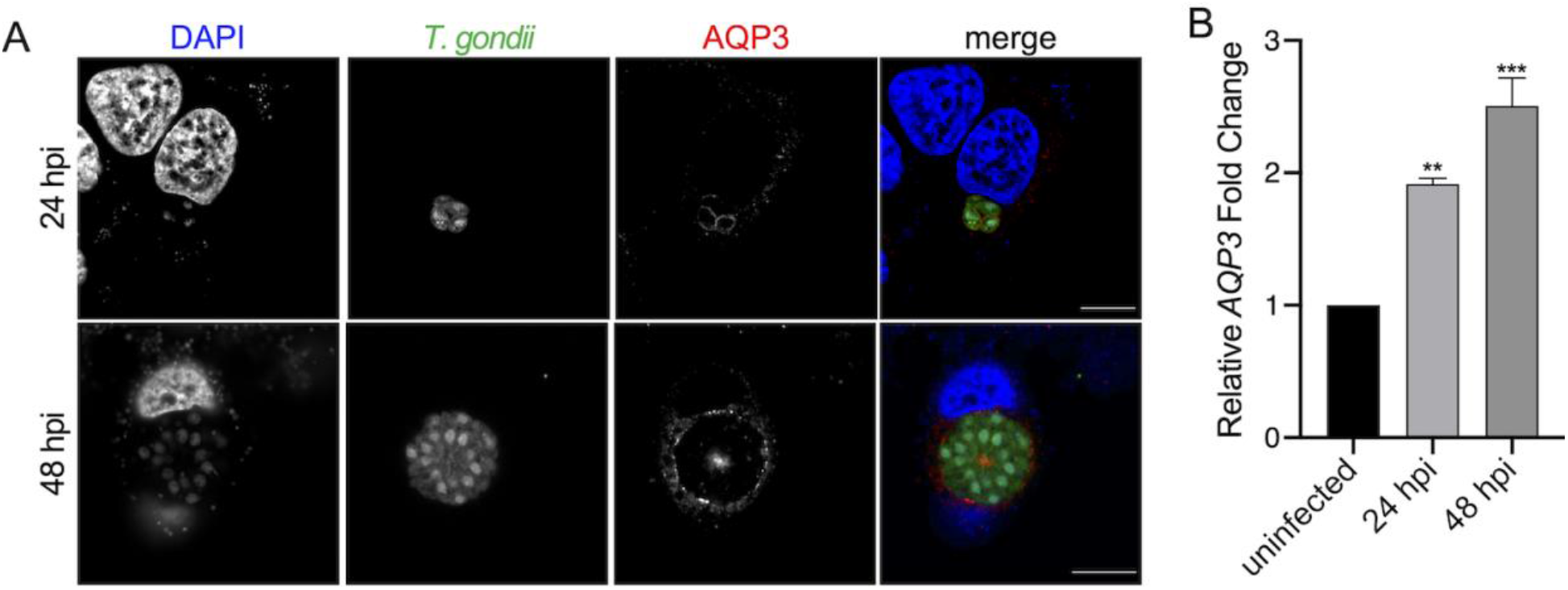
AQP3 recruitment and upregulation during *T. gondii* infection. (A) Representative confocal images of HuH7 cells infected with GFP-expressing *T. gondii* Pru A7 (green), stained with DAPI (blue) and anti-*Hs*AQP3 (red) at 24 and 48 hpi. Scale bar is 10 µm. (B) qRT-PCR quantification of host *AQP3* mRNA expression in uninfected and *T. gondii* Pru A7-infected HuH7 cells at 24 and 48 hpi. Infected cells at 24 and 48 hpi exhibit a 1.9 ± 0.043-fold (*p*=0.0034) and 2.5 ± 0.21-fold (*p*=0.0003) increase, respectively, in *AQP3* expression (Dunnett’s multiple comparisons test; n=3 independent experiments). Data reported as mean ± SEM; n=3 independent experiments. \*\**p* < 0.01, \*\*\**p* < 0.001.

Based on the relative increase in AQP3 fluorescence signal observed in our confocal images of uninfected versus *T. gondii*-infected cells, we predicted gene expression was upregulated after infection, similar to the upregulation observed after *P. berghei* infection of hepatocytes (Posfai et al., 2018). To test this, we evaluated *AQP3* levels in HuH7 after infection with *T. gondii* Pru A7. In these experiments, we typically observe ∼50% parasite infection rates, therefore cells were not sorted but analyzed as a mixed population of infected and uninfected cells. We observed that *AQP3* is upregulated at 24 (1.9-fold change, *p*=0.0034) and 48 (2.5-fold change, *p*=0.0003) hpi when compared to uninfected cells (Figure 4B). We also observed an increase in *AQP3* expression in Vero cells infected with *T. gondii* Pru A7 (Figure 4—figure supplement 2C) and HuH7 cells infected with *T. gondii* RH (Figure 4—figure supplement 2D), demonstrating the phenotype in both type I and II strains, and with various host cells.

To examine the importance of AQP3 in *T. gondii* infection, we employed a chemical biology approach and examined parasite growth in the presence of the known AQP3-inhibitor auphen (Martins et al., 2012). To complete this study, we first optimized the infection of HuH7 cells by a *T. gondii* Me49-luciferase strain (Pernas et al., 2014) in 384-well microplates for high-throughput screening (Z-factor=0.82). In this assay, compounds are added at 0 hpi and then parasite load concurrent with host viability is assessed at 48 hpi. Using this method, we observed that auphen inhibits *T. gondii* Me49 parasite load in HuH7 cells with an EC_50_ value of 2.3 ± 0.61 µM. At auphen concentrations above 10 µM, the near-complete inhibition of parasite load (100% reduction) was observed (Figure 5A). We also examined the change in *T. gondii* Pru A7 rosette size in HuH7 cells as a function of auphen concentration (Figure 5B). This analysis showed that auphen reduces rosette size with an EC_50_ value of 1.8 ± 0.47 µM. At high auphen concentrations (>10 µM), rosette size decreased (Figure 5C) by >90% when assessed at 48 hpi. No change was observed in the *T. gondii* Pru A7 infection rate in the presence of auphen (50 or 100 µM) when compared to DMSO at 24 hpi (Figure 5—figure supplement 1A). A similar auphen potency was determined for reduction of *T. gondii* Pru A7 parasite size in Vero cells (EC_50_=1.8 ± 0.14 µM). Additionally, we determined that *T. gondii* Pru A7 rosette size in Vero cells was reduced 71% in the presence of 10 µM auphen (Figure 5—figure supplement 1B). Our observed auphen inhibition of *T. gondii* is similar to the potency previously reported against various *Plasmodium* species and stages (Figure 5D).

**Figure 5.**
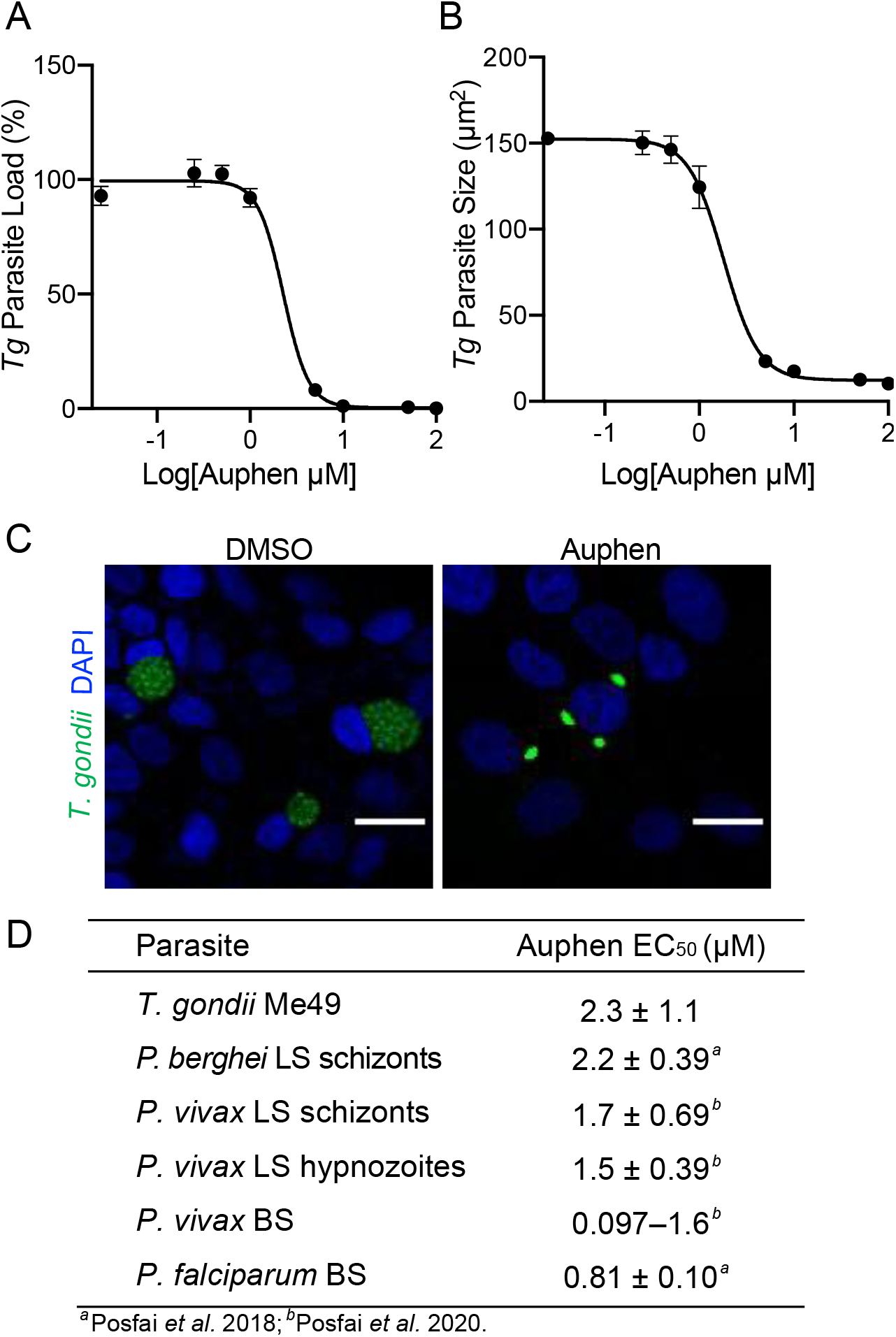
Auphen inhibition of apicomplexan parasites. (A) Auphen dose-dependent inhibition of luciferase-expressing *T. gondii* Me49 in HuH7 cells assessed at 48 hpi, evaluated with relative luminescence. Auphen EC_50_ = 2.3 ± 1.1 µM (mean ± SEM; n=3 independent experiments). (B) Auphen dose-dependent decrease of GFP-expressing *T. gondii* Pru A7 rosette size in HuH7 cells assessed at 48 hpi, evaluated with confocal microscopy and analyzed with ImageJ. Auphen EC^50^ = 1.8 ± 0.47 µM (mean ± SEM; n=2 independent experiments). (C) Representative confocal images (20x) of GFP-expressing *T. gondii* Pru A7 in HuH7 cells in the presence of DMSO (left) or 50 µM auphen (right) at 48 hpi. Images display *T. gondii* Pru A7 (green) and DAPI (blue). Scale bar is 50 µm. (D) Table with EC^50^ values for auphen inhibition of *T. gondii* Me49 parasite load in HuH7 cells and various *Plasmodium* parasites. Parasite species and stage (LS, liver-stage; BS, blood stage) indicated for *Plasmodium*. Data shown as mean ± SEM, except for *P. vivax* BS which is the range determined from various isolates.

## Discussion

*P. vivax* remains a critical hurdle to malaria eradication efforts due in large part to the parasite’s ability to persist in the liver and lead to relapses in blood infections. During the liver stage, the parasites can mature into schizonts after invasion or delay replication as a seemingly arrested form termed a hypnozoite, which can unpredictively activate. Processes that drive invasion, maturation and development during this time are considered elusive, yet much less is understood about contributing factors that enable the survival of dormant hypnozoites. Our study provides insights into molecular processes occurring during this liver stage by highlighting the role of the TVN in both *P. vivax* schizonts and hypnozoites. Previously well-characterized in liver-stage *P. berghei* schizonts, the TVN is known to be critical for nutrient acquisition and evasion of the host immune responses (Agop-Nersesian et al., 2018, 2017; Thieleke-Matos et al., 2016).

Our study provides the first description of the TVN in *P. vivax* EEFs. Interestingly, we observed differences in the utilization of this network throughout infection between *P. vivax* and *P. berghei*. Both *Plasmodium* species exhibit a similar sequence of developmental events that lead to schizonts, which includes invasion, morphological changes, replication, merozoite formation, and parasite release, though at varying time scales. Based on a previous report, *P. berghei* exhibits greater TVN activity before replication (Grützke et al., 2014), at early time points, yet we observe this network is present throughout a 10-day *P. vivax* time course. The rise in TVN activity between 2-6 dpi highlights an important functional role during that time. Conversely, the decline observed in TVN features after 6 dpi likely signifies a change in the EEF development, whereby TVN function is less critical. At these later stages of development, TVN-derived vesicles are the most prominent feature. TVN vesicles are expelled into the host cytosol and Grützke and colleagues have reported that they have a relatively slower velocity compared to the other studied features (Grützke et al., 2014). One possibility for our observation of these features after others have dissipated is that they may take longer to recycle but their presence throughout liver-stage infection suggests this is unlikely due to ample time for clearance. Another possibility is that vesicles observed at 9 and 10 dpi in schizonts may facilitate egress or further protect the parasite from clearance by shedding autophagy related proteins to these TVN-derived vesicles. A real-time evaluation of TVN characteristics would be critical for evaluating the dynamics of each feature and correlating them with developmental changes within the parasites and protection from clearance by the host. This remains difficult as the genetic tools necessary to generate transgenic *P. vivax* sporozoites that constitutively express UIS4 are not yet available. Such a tool would also be critical to assess TVN features that may have been missed due to the time points selected for microscopy analysis.

While the TVN in developing *P. vivax* EEFs was anticipated based on *P. berghei* studies, its presence in hypnozoites, which are considered biologically quiescent, was less expected. In fact, at later days when hypnozoites and liver-stage schizonts can be distinguished microscopically, TVN features were more prevalent in *P. vivax* hypnozoites versus schizonts. This host-pathogen activity seems counterintuitive to notions of the dormant state and suggests that many as yet unknown complex interactions occur during this time. Indeed, recent RNA-sequencing studies of *P. vivax* hypnozoites have suggested metabolic activity occurs at this time (Gural et al., 2018). In regard to the TVN, one explanation for these features is that they prime the dormant EEF for activation, essentially preparing the parasite for nuclear division. We observe a significant decrease in hypnozoites from day 6 (67%) to day 10 (19%), suggesting activation, or alternatively host clearance, may occur during this time. Although hypnozoite activation is an enticing hypothesis for TVN activity, the lack of knowledge about this process makes it difficult to prove. It is also possible that the EEFs displaying these features are able to evade the host immune response, something that may be further explored with future studies evaluating the association of host immune response proteins to the TVN present in hypnozoites.

We observed host AQP3, a protein channel that is permeable to water, glycerol and other small solutes, associates with the TVN in both *P. berghei* and *P. vivax* EEFs. Interestingly, various host aquaporins have been implicated in apicomplexan infection. Previous studies have shown *Cryptosporidium parvum* invasion is aided by aquaporin 1 and localizes to the host–parasite interface (Chen et al., 2005). Further, an RNA-seq study demonstrated the upregulation of *AQP3* and *AQP4* in mice with chronic-infection of *T. gondii* (Pittman et al., 2014). The present study adds to this possible association by demonstrating an increase in *AQP3* levels after *T. gondii* infection of hepatocytes and a reduced size phenotype for *T. gondii* in auphen-treated HuH7 cells, similar to that previously observed with *P. berghei* (Posfai et al., 2018). Thus, host aquaporins may represent a class of host proteins with significance to apicomplexan infection. This function may be complementary to that of apicomplexan aquaporins, which can have essential functions in transporting water and small solutes (Hansen et al., 2002; Miranda et al., 2010; Promeneur et al., 2018, 2007).

A well-characterized role for AQP3 in mammalian cells is in nutrient flux, a function similar to that of the TVN in liver-stage *P. berghei* EEFs (Agop-Nersesian et al., 2018). We detected AQP3 exclusively in extended membrane clusters and TVN-derived vesicles in *P. vivax* EEFs throughout infection (Figure 3B), suggesting important and coincident functions. Notably, we did not observe localization of AQP3 to all extended membrane clusters and vesicles. This partial co-localization was expected due to the transitionary state of AQP3 between 2–5 dpi, where the protein is being recruited to the *P. vivax* PVM (Posfai et al., 2020). It remains unknown if AQP3 is trafficked from the host cytosol to the PVM by TVN-derived vesicles or alternatively, AQP3 in the PVM is shed into vesicles. In support of the former hypothesis, we did not observe AQP3 localization in TVN-derived vesicles without localization to PVM concurrently. Among the host proteins known to associate with the liver-stage TVN, it has been reported that both LC3 and LAMP1 are shed from the PVM to the TVN to facilitate evasion of the host immune response (Agop-Nersesian et al., 2017; Thieleke-Matos et al., 2016). While this highlights a role for the TVN in protection from the host response, the network is also known to be important for nutrient acquisition. Previous reports have shown that host-derived endosomes and lysosomes associate with the TVN, presumably releasing their degraded contents to the network where they can then potentially be imported as nutrients (Agop-Nersesian et al., 2017; Grützke et al., 2014). AQP3 could therefore facilitate this nutrient acquisition. A possible function in nutrient acquisition may not be relevant to tubules, which may account for the lack of AQP3 association with that TVN feature. It also remains possible that we could not detect AQP3 association to TVN tubules due to their highly dynamic nature (Grützke et al., 2014) and our method of analysis, but we were confidently able to establish the connection between the host protein and TVN-extended membrane clusters and vesicles. AQP3 has also been associated with oxidative stress and the immune response, often via its ability to transport hydrogen peroxide (Hara-Chikuma et al., 2015; Miller et al., 2010; Nalle et al., 2020; Vieceli Dalla Sega et al., 2014; Wang et al., 2020). Thus, hydrogen peroxide or other as yet unidentified solutes remain plausible functions for AQP3 in the PVM/TVN of *P. vivax* schizonts and hypnozoites.

Beyond our efforts with *Plasmodium*, we were able to demonstrate AQP3 is recruited to *T. gondii*, a model apicomplexan parasite. Specifically, we observed *AQP3* upregulation and recruitment to the PPM at 24 hpi and to the PVM at 48 hpi in *T. gondii* Pru A7-infected hepatocytes. Interestingly, *AQP3* induction was also observed in chronic *T. gondii*-infected mice (Pittman et al., 2014). At 24 hpi, we observed AQP3 recruitment to the PPM in ∼10% of *T. gondii-*infected HuH7 cells. This recruitment changed by 48 hpi where AQP3 was observed in the PVM in ∼50% of infected cells. At this time, we also observed AQP3 localization to the residual body. While not present in *Plasmodium*, the residual body is known to help structural components of the rosette, facilitate division, and exchange solutes in *Toxoplasma* (Coppens and Romano, 2018; Frénal et al., 2017; Periz et al., 2017). Therefore, the presence of AQP3 at this junction could aid in solute transport. Notably, *T. gondii* also displays a TVN, although this membranous structure is present between the PPM and the PVM, whereas in the *Plasmodium* LS, the TVN is comprised between the PVM and host cytosol. While we were unable to detect the *T. gondii* TVN under our experimental conditions, the recruitment of AQP3 suggests that it may aid in survival for some parasite populations.

Of note, only ∼50% of *T. gondii* rosettes have AQP3 recruitment by 48 hpi when compared the 100% recruitment observed with *P. berghei* EEFs. Counterintuitive to this observation, we found that the known AQP3 inhibitor auphen completely inhibited *T. gondii* load in HuH7 cells. This finding suggests that additional parasite and/or host targets exist for auphen as it inhibits all parasites regardless of AQP3 recruitment. Examining single-cell gene expression in cells without AQP3 recruitment may shed light on factors that can compensate for its function during parasite infection. Importantly, *AQP3* gene induction, AQP3 protein recruitment and parasite sensitivity to auphen were observed in three different *T. gondii* strains (Pru A7, Me49 and RH), from two lineages (type I and type II), and with different host cells (HuH7 and Vero). These results point to *T. gondii* as a possible model for future studies investigating AQP3 function and inhibitor screening.

Taken together, we demonstrate a previously unknown role for the TVN in dormant *P. vivax* hypnozoites and highlight the possible function of AQP3 in this network in both mouse- and human-infective liver-stage *Plasmodium*. AQP3 association specifically with TVN-extended membrane clusters and vesicles implicates the water and solute channel in their function throughout infection. Further, our demonstration that AQP3 is also recruited to *T. gondii* suggests a broad importance of this protein to apicomplexans. Future efforts to resolve the molecular function of AQP3 throughout schizont and hypnozoite development as well as studies to map host and parasite constituents of the TVN will be critical to advancing our understanding of host-parasite interactions during this critical time of the life cycle.

## Materials and Methods

### Lead contact and materials availability

Further information and requests for *P. vivax-*related reagents should be directed to BenoÎt Witkowski. There are restrictions on the availability of some *P*. vivax-related reagents due to inadequate methodology for the preservation and propagation of the parasites in clinical isolates. This study did not generate new unique reagents from Emily Derbyshire. Further information about auphen and imaging reagents should be directed to and will be fulfilled by the Lead Contact, Emily Derbyshire.

### *P. vivax* infections of primary human hepatocytes (PHH)

*P. vivax* infections were completed as previously described (Roth et al., 2018). Blood samples were collected from symptomatic *P. vivax* patients at local health facilities in Mondulkiri province (eastern Cambodia) from 2018-2019. Clinical isolate collection and research procedures were reviewed and approved by the Cambodian National Ethics Committee for Health Research (approval number: #101NECHR& #273NECHR). Patients presenting signs of severe malaria, infected with non-vivax malaria parasites, under 5 years of age, or who were pregnant or lactating were excluded from the collection. Following informed consent from eligible study participants, venous blood samples were collected by venipuncture into heparin-containing tubes (Beckton Dickinson, Cat# 367886). Venous blood was pelleted and serum was replaced with naïve human serum (Interstate Blood bank, Inc.) before feeding to *An. dirus* mosquitoes using glass bell insect feeders held at 37°C. Following a *P. vivax* gametocyte-containing bloodmeal, *An. dirus* mosquitoes were maintained on a natural light cycle and 10% sucrose in water. Mosquitoes found positive for *P. vivax* oocysts at six days post-feeding were transported to the IPC facility in Phnom Penh, Cambodia where salivary glands were aseptically dissected into RPMI without sodium bicarbonate (Gibco, Cat# 61870-010) on day 16-21 post-feeding. Cryopreserved PHH were thawed into InVitroGro™ CP Medium (BioIVT) including a 1x antibiotic mixture (PSN, Gibco and Gentamicin, Gibco) and 18,000 live cells were added to selected wells of a collagen-coated 384-well plate (Grenier). Cultures were maintained in a standard tissue culture incubator at 37 °C and 5% CO_2_. Two lots of PHH were used for IFA analysis: lot BGW was obtained from a 50-year old Caucasian male, lot UBV was obtained from a 57-year old Caucasian male. Infection of PHH was performed at 2 days post-seed by diluting freshly dissected sporozoites into CP media with antibiotics, adding 20 μL sporozoite-media mixture to each well, and centrifugation the 384-well plate at 200 RCF for 5 min at room temperature. Media was exchanged with fresh CP media containing antibiotics the day after infection and every 2-3 days thereafter.

### Liver stage *P. vivax* immunofluorescent (IFA) microscopy and TVN analysis

*P. vivax*-infected PHH were fixed with 4% paraformaldehyde (ThermoFisher) for 1 hr. Fixed cells were stained with recombinant mouse anti-*P. vivax* Upregulated in Infectious Sporozoites-4 antibody (rPvUIS4, (Schafer et al., 2018)) (1:2,500) in staining buffer (0.03% TritonX-100, 1% (w/v) BSA in PBS) overnight at 4°C. Cells were then washed with PBS and stained with rabbit anti-mouse Alex Fluor® 488-conjugated antibody (1:1000) in staining buffer overnight at 4 °C. Cells were then washed with PBS, stained with rabbit anti-*Hs*AQP3 (AbClonal) (1:200) for 48 hrs at 4°C, washed again with PBS and incubated with donkey anti-rabbit Alexa Fluor® 568-conjugated antibody (ThermoFisher) (1:400). Stained cells were washed with PBS and counterstained with 0.5 μg/mL DAPI (ThermoFisher). Fluorescence was detected using a Zeiss 880 Airyscan inverted confocal and images were analyzed using ImageJ (Schindelin et al., 2012). Infections were evaluated as 2D and Z-stacked images to assess TVN structures present. AQP3 colocalization with TVN structures was observed throughout *P. vivax* liver-stage infection (2– 10dpi). GraphPad Prism was used to generate quantitative figures.

### *P. berghei* infections of HuH7 cells

HuH7 cells (kind gift from Dr. Peter Sorger) were cultured in DMEM + L-Glutamine (Gibco) supplemented with 10% (v/v) heat-inactivated FBS (HIFBS) and 1% (v/v) antibiotic/antimycotic (Sigma). Cultures were maintained in a standard tissue culture incubator at 37°C and 5% CO_2_. *Anopheles stephensi* mosquitoes infected with luciferase-expressing *P. berghei* ANKA were purchased from NYU Langone Medical Center Insectary. HuH7 cells were seeded (2.5×10^5^/well) into a 24-well plate with coverslips 24 hours prior to infection. Cells were infected with ∼50,000 *P. berghei* sporozoites per well that were freshly dissected from *An. stephensi* mosquitoes.

### Liver-stage *P. berghei* IFA microscopy and TVN analysis

*P. berghei*-infected HuH7 cells were fixed at 24 hpi or 48 hpi with 3% paraformaldehyde at room temperature for 15 min. Cells were permeabilized with 0.2% TritonX for 10 min, washed with PBS and blocked with 3% BSA for 1 hr at room temperature. Cells were stained with goat anti-*Pb*UIS4 (antibodies-online) (1:1000) for 1 hr at room temperature, washed with PBS and incubated with secondary donkey anti-goat Alexa Fluro® 488-conjugated antibody (ThermoFisher) (1:400). Cells were then stained sequentially with rabbit anti-AQP3 (Rockland) (1:100) overnight at 4°C and donkey anti-rabbit (ThermoFisher) (1:400) for 1 hr at room temperature. Lastly, cells were stained with DAPI. Fluorescence was detected using a Zeiss Axio Observer Widefield Fluorescence Microscope and images were analyzed using ImageJ.

### *T. gondii* infections of host cells

Vero (ATCC) and HeLa (ATCC) cells were cultured in DMEM with L-glutamine (Gibco) supplemented with 10% (v/v) HI-FBS and 1% (v/v) antibiotic/antimycotic (Sigma) in a standard tissue culture incubator at 37 °C with 5% CO_2_. *T. gondii* tachyzoites of the type II strain Prugniaud (Pru) A7 (a kind gift from Dr. Jörn Coers) and luciferase-expressing ME49 (a kind gift from Dr. John Boothroyd) were propagated in Vero or HeLa cells. Type I mCherry-expressing RH *T. gondii* (a kind gift from Dr. Laura Knolls) were propagated in HeLa cells. Vero or HeLa cells infected with parasites were scraped, and host cells were lysed by passing them through a 25-gauge needle six times to release tachyzoites. For microscopy, Vero cells (2.5 x 10^5^) were seeded on coverslips in 24-wells plates and infected with *T. gondii* with a multiplicity of infection (MOI) of 5.

### *T. gondii* IFA microscopy and analysis

*T. gondii* Pru A7-infected HuH7 cells were fixed at 24 hpi or 48 hpi with 4% paraformaldehyde for 15 min at room temperature. Cells were washed with PBS and blocked with 3% BSA for 1 hr at room temperature. Cells were incubated with rabbit anti-*Hs*AQP3 (AbClonal) (1:200) for 72 hours at 4 °C, washed with PBS and incubated with donkey anti-goat Alexa Fluor® 568 (ThermoFisher) (1:400) for 1 hr at room temperature. After washing with PBS, cells were stained with DAPI (0.5 µg/mL) and washed once more. ProLong® Gold Antifade (ThermoFisher) was added to stained wells for preservation of fluorescence. Fluorescence was detected using a Zeiss 880 Airyscan inverted confocal microscope and images were analyzed using ImageJ. A minimum of three independent experiments, 50 parasites/experiment, were completed for all localization studies.

*T. gondii* parasite size in the presence of auphen was assessed with high-content imaging in Vero cells. Auphen (20–0.01 µM) was added to *T. gondii* Pru A7-infected Vero cells before infection. Vero cells (4,000 cells/ wells) were infected with a MOI of 0.375 in a 384-well plate. All wells were at a final volume of 30 uL and a final concentration of 1% DMSO. The number of rosettes in every DMSO- and auphen-treated well was quantified 48 hpi. *T. gondii* parasite size was assessed using the GFP reporter and DAPI was used to assess Vero cell numbers using the Cellomics ArrayScan (ThermoFisher) high-content imaging system. EC_50_ values were determined by fitting data to a standard dose-response equation (GraphPad Prism). Two independent experiments were performed with 5-10 wells/condition in each experiment.

### qRT-PCR

*T. gondii*-infected HuH7 cells were harvested from 6-well plates 2 dpi and resuspended into lysis buffer. RNA was extracted using the Quick-RNA miniprep (Zymo), and cDNA was synthesized using GoScript™ Reverse Transcriptase System (Promega) according to the manufacturer’s instructions. The qRT-PCR analysis was performed with the oligonucleotide primers listed in STAR Methods and SYBR Green I Master reagents with a LightCycler® 480 Instrument II (Roche Diagnostics). The final volume was 5 µL in a 384-well plate. *Hs*AQP3 levels were normalized to *H. sapiens* 18S in *T. gondii*-infected and uninfected cells. Each sample was analyzed in triplicate from 3 independent experiments.

### *Toxoplasma* parasite inhibition assays

HuH7 cells (7,000/well) were seeded in 384-well plates in the presence or absence of auphen (20–0.01 µM) in triplicate before infection with luciferase-expressing *T. gondii*-Me49 tachyzoites (6,000 parasites/well). After 45 hrs, HuH7 viability and *T. gondii* parasite load were assessed using the protocol described above. The relative signal intensity of each well was normalized to the negative control (1% DMSO). The positive control was pyrimethamine (10 µM). EC_50_s were determined using GraphPad Prism and the reported values are the average determined from three independent experiments.

## Acknowledgments

We thank the New York University and the University of Georgia, Athens Insectaries for providing *P. berghei-*infected mosquitoes, the Duke Microscopy Core Facility and the Duke Functional Genomics Facility. We thank Dr. Jörn Coers (Duke Medical School), Dr. Laura Knolls (UW-Madison) and Dr. John Boothroyd (Stanford University) for providing *T. gondii* strains, Prugniaud A7 (GFP), RH (mCherry), and ME49 (luciferase), respectively. We thank the *P. vivax* patients of Mondulkiri Province, Cambodia for participating in this study.The recombinant mouse anti-*Pv*UIS4 antibody was obtained from Noah Sather of Seattle Children’s. We also thank Jack G. Ganley for thoughtful discussions on this paper and providing auphen. Work was supported by the Bill & Melinda Gates Foundation (OPP1023643 D.E.K.), the NIH (1DP2AI138239 to E.R.D) for laboratory support, and fellowship support by the NSF (DGE-1644868, to K.S.).

## Footnotes

The authors have no competing interests.

## Notes

### Competing Interest Statement

The authors have declared no competing interest.

